# Pooling in a predictive model of V1 explains functional and structural diversity across species

**DOI:** 10.1101/2021.04.19.440444

**Authors:** Angelo Franciosini, Victor Boutin, Frédéric Chavane, Laurent U Perrinet

## Abstract

Neurons in the primary visual cortex are selective to orientation with various degrees of selectivity to the spatial phase, from high selectivity in simple cells to low selectivity in complex cells. Various computational models have suggested a possible link between the presence of phase invariant cells and the existence of cortical orientation maps in higher mammals’ V1. These models, however, do not explain the emergence of complex cells in animals that do not show orientation maps. In this study, we build a model of V1 based on a convolutional network called Sparse Deep Predictive Coding (SDPC) and show that a single computational mechanism, pooling, allows the SDPC model to account for the emergence of complex cells as well as cortical orientation maps in V1, as observed in distinct species of mammals. By using different pooling functions, our model developed complex cells in networks that exhibit orientation maps (e.g., like in carnivores and primates) or not (e.g., rodents and lagomorphs). The SDPC can therefore be viewed as a unifying framework that explains the diversity of structural and functional phenomena observed in V1. In particular, we show that orientation maps emerge naturally as the most cost-efficient structure to generate complex cells under the predictive coding principle.

**Significance:** Cortical orientation maps are among the most fascinating structures observed in higher mammals brains: In such maps, similar orientations in the input image activate neighboring cells in the cortical surface. However, the computational advantage brought by these structures remains unclear, as some species (rodents and lagomorphs) completely lack orientation maps. In this study, we introduce a computational model that links the presence of orientation maps to a class of nonlinear neurons called complex cells. In particular, we propose that the presence or absence orientation maps correspond to different strategies employed by different species to generate invariance to complex stimuli.

## 1 Introduction

Cells in the primary visual cortex of higher mammals (V1) have historically been divided into two classes: simple and complex [1]. Simple cells show linear responses to drifting sinusoidal grating of a specific frequency. In particular, it has been shown that the simple cells’ responses are modulated by drifting gratings and are thus maximal when the stimulus matches a specific phase inside their receptive field (RF) [1–3]. On the other hand, complex cells show the same selectivity for oriented patterns but, in contrast, their responses are not modulated by the drifting phase and remain almost constant throughout the whole cycle [1, 2, 4, 5]. As such, complex cells are thought to be encoding for edges of a specific orientation independently from their phase or position [6]. The presence of these two different cell types is in line with the idea that the brain solves the challenging task of object recognition by decomposing it into a series of simpler problems along a computational hierarchy [7]. In fact, the emergence of complex cells in V1 can be explained by hierarchical models where simple cells feed information into complex cells through a nonlinear spatial pooling [8, 9]. This hypothesis is consistent with the remarkable hierarchical structure of mammal’s visual system: Neurons respond to more complex stimuli moving along the ventral stream: from simple cells responding to simple edges in the primary visual cortex (V1), shapes and textures in V4 [10] and specific objects in the inferotemporal region (IT) [11]. In this computational chain, complex cells are thought to play a role in building invariance to the stimulus’ properties within V1 and for the sensory information to be processed upstream in the hierarchy [12]. Such a mechanism could implement an efficient algorithm for the detection of the objects’ boundaries and indeed, it represents a crucial computational step in numerous models of contour integration and object recognition of human vision [7, 13–15]. How the brain develops this structure, however, remains an open question.

One successful model in understanding the responses of V1 cells is sparse coding (SC) [16]. This framework has long been used to model the strategy employed by mammals’ primary visual cortex (V1) to detect low-level features. According to SC, neurons in V1 try to reconstruct the input signal while attempting to respond in the most energy-efficient way possible, generating a sparse activity pattern where the majority of cells are inactive and only a few are firing. Sparse activity is in line with the efficient coding hypothesis stated by Barlow in 1961 [17]: Neural systems actively minimize the total firing required to represent sensory information. SC can achieve this by refining the neural responses through a recurrent inference process. Interestingly, when trained on natural images, sparse coding can model the diversity of the receptive fields (RFs) properties found in the primary visual cortex of macaques [18]. Although this method is efficient in modeling simple (shallow) visual networks and the emergence of simple cells, sparse coding does not directly tackle deeper hierarchical processing and, in general, fails to predict the emergence of complex cells. In particular, extending its mathematical framework to account for visual hierarchies is nontrivial. SC also fails to predict the emergence of extra-classical receptive field effects, that is, the ability of a stimulus presented outside its classical RF to modulate the response of a neuron to a stimulus inside its classical RF.

One theory that explicitly addresses the problem of hierarchical processing in the brain is Predictive Coding (PC). This framework introduced by Rao & Ballard [19] exploits feedback and feed-forward connectivity to solve a Bayesian inference problem by modeling visual processing as a cascade of dynamical systems organized along the hierarchy of the visual pathways [20]. Predictive Coding suggests that every cortical area predicts at best the upstream sensory information. The mismatch between the prediction and the lower-level activity elicits a prediction error that is used to adjust the neural response until an equilibrium is reached. Information transfer is made possible by feedback and feed-forward connectivity; the first carries predictions while the second transmits prediction errors [21]. With this simple idea, PC offers on one hand an elegant explanation for the abundance of feedback connectivity in the brain [22] and on the other hand, extra-classical RF effects in V1 [19].

To take advantage of the modeling power of SC and PC we integrated both these mechanisms in a model of V1, called Sparse Deep Predictive Coding (SDPC). This framework introduces biologically realistic convolutional architecture which integrates predictive coding while accounting for sparsity constraints [23, 24]. Yet, both these SC and PC models are missing a third key component necessary to model the emergence of complex cells: nonlinear pooling. Sakai and Tanaka [8] showed that classical cascade models fail to predict phase-invariance. The existence of complex cells in V1 can be explained by hierarchical models where simple cells feed information into complex cells through a nonlinear spatial pooling, which is usually modeled as the sum of squared simple cells responses or a winner-take-all mechanism (also known as max-pooling) [25]. Thanks to pooling, a single complex cell could respond to the same stimulus, independently of its exact position, efficiently generating invariance when stimulated with a drifting grating. Indeed, visual areas in mammals share the common property of being organized in *retinotopic maps* [26,27]: cells on the cortical surface can be mapped to contiguous locations on the retinal surface. In other words, neighboring cells in the visual cortex respond to neighboring regions of the visual field, defining a spatial encoding within the cortical space. We predict that complex cells could emerge by pooling across neighboring positions in the retinotopic map, given a cortical pooling region large enough to span across different spatial locations.

Another structure of the cortex that could explain the emergence of complex cells is the presence of an *orientation map*, that is, a functional structure of the selectivity of cells for which neighboring cells on the cortical surface are tuned to similar orientations. Orientation preference varies smoothly but also shows local singularities, called pinwheels [27–29]. Similar to the retinotopic feature, pooling across the response of local groups of neurons in this topological structure could explain the emergence of phase invariance in complex cells, while keeping selectivity to orientation. Important results in this sense were obtained by Hyvarinen *et al.* [30], by defining a sparse coding model that included pooling that efficiently reproduces complex cells’ responses in V1. In addition, such a model offers a functional link between complex cells and topographic orientation maps observed in higher mammals’ visual cortex. Similar results were obtained with a type of independent component analysis (ICA) [31]. Remarkably, Antolik *et al.* [9] managed to build a neuro-inspired model that extended the principle of topographic maps to complex cells. It is important to note at this point that neurophysiological evidence shows that there is no evident spatial structure or orientation preference in rodents (most notably mice, rats, and squirrels), although single neurons are observed to be orientation selective [27, 29]. This type of organization is commonly referred to as salt-and-pepper. Besides, these species, while not having clear orientation maps, do have V1 cells which show complex cell properties. The models listed above, however, cannot explain the presence of these complex cells in animals that do not present topographic maps in V1. Altogether, these observations open two important questions: *How are orientation maps and complex cells related? Can the same framework be used to explain complex cells emergence in animals that lack orientation map?*

Here, to shed light on the link between cortical maps and complex cells, we propose to include sparse, hierarchical processing, and nonlinear pooling within a single model of the primary visual cortex. In this study, we show that the SDPC can develop both complex cells and orientation maps, as well as complex cells without orientation maps, and we interpret these results under the predictive coding principle. Interestingly, we found that a single mechanism (pooling) is sufficient to explain different cortical architectures across species. We define two distinct pooling functions that act in different subspaces: the spatial retinotopic space, and a feature space that can be learned by the network in an unsupervised fashion. First, we show that biologically plausible orientation maps emerge naturally in the feature space as the most cost-efficient structure, given the right pooling function. Second, we show that these orientation maps represent the most cost-efficient structure that allows the emergence of complex cells. These results allow us to present SDPC as a unifying model of the early visual cortex, which building principles can explain the link between topographical structures (retinotopic maps and orientation maps) and functional phenomena (complex cells). The paper is organized as follows: First, we describe our model by detailing its computations. In particular, we will concentrate on briefly outlining the SDPC model and introduce the pooling functions used to integrate local information. Then, we will present simulations of this model in light of current neuroscientific knowledge. Finally, we will discuss the implications of this work with respect to the state-of-the-art.

## 2 Methods

### 2.1 The SDPC framework

The Sparse Deep Predictive Coding (SDPC) framework solves a series of hierarchical inverse problems with sparsity constraints. A group of neurons *γ_i_* predicts, at best, the activity from the previous cortical layer *γ_i_*_−1_, trough a set of synaptic weights *W_i_*. Given a network with *N* layers, we can define the generative model [23, 24] as:

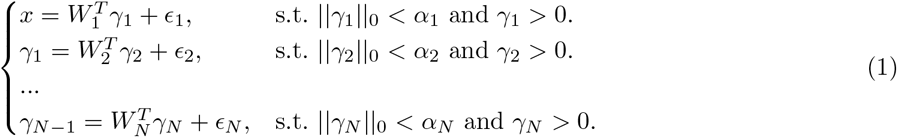

Where *x* represents the input stimulus and *γ_i_* are the rate-based neural responses for layer *i* (see Fig. 1, b). Sparsity constraints are introduced using the *ℓ*_0_ pseudo-norm which computes the number of active elements in each activity map *γ_i_*. The *W_i_* matrices represent the network’s weights (convolutional kernels) which will be described later, and *E_i_* is the prediction error associated to each layer. One limit of this formulation is that it only allows the stacking of sparse, yet linear, inverse problems. In order to model complex cells, we wish to introduce an additional nonlinear pooling defined as functions *p_i_*(*γ_i_*) (Fig. 1, a). We modify the generative problem as follows:

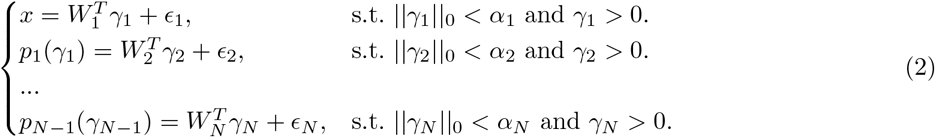

**Fig 1.**
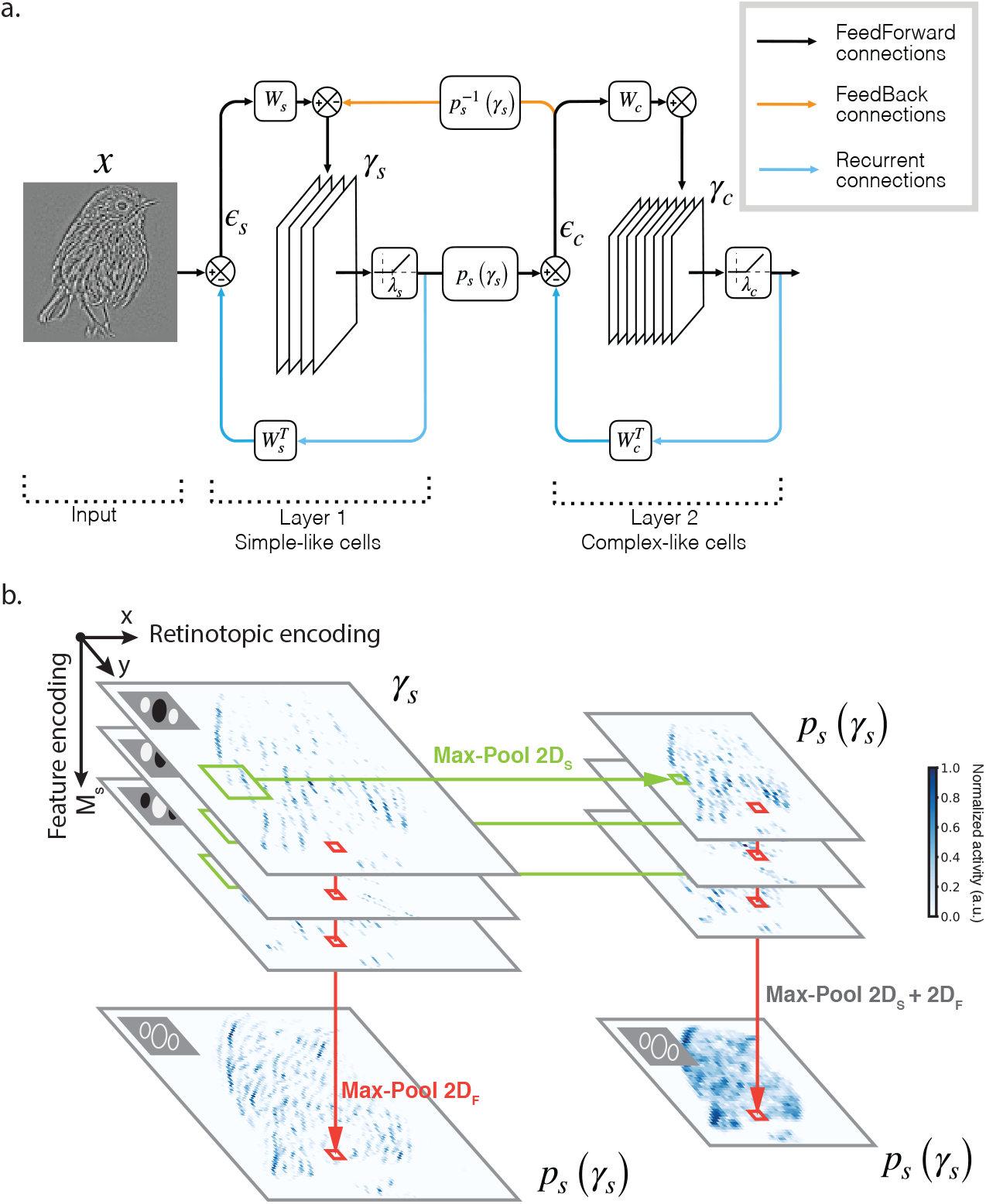
The SDPC network. **a.** Update scheme of the SDPC network used in this study: *x* is the input image, *γ_s_* and *γ_c_* represent simple and complex cells response maps, respectively. *W_s_* and *W_c_* are convolutional kernels that encode for the RFs of the simple and complex cell layers, where each synaptic weight matrix is composed of *M_s_* and *M_c_* neurons (kernels) respectively. *p_s_*(*γ_s_*) is the pooling function used to generate position and feature invariance in the response of the second layer. feed-forward connections carry information on the prediction errors (*ϵ_s_* and *ϵ_c_*) that is used to refine the neural activities. Feedback information is carried through the unpooling function 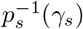, that approximates the derivative of the pooling function. **b.** Here, we show a representation of *γ_s_* and *p* (*γ_s_*). Each pixel represents a model neuron and the color code indicates the amplitude of the neural response (lighter for no response and darker blue for the maximal response, here normalized to 1). In this figure, we illustrate three possible outputs *p_s_* (*γ_s_*), used to generate different network structures: MaxPool 2*D_S_* selects the maximum activity over spatial (retinotopic) positions, in each plane independently; MaxPool 2*D_F_* acts in the feature space by selecting the maximum activity across planes in *γ_s_*. When the network integrates the two pooling functions in sequence, we refer to them as a unique max-pooling operator called MaxPool 2*D_S_* + 2*D_F_*.

The optimal state of the neural activity for each layer can be found by defining a series of local loss functions:

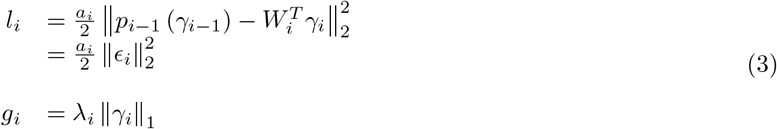

Where *l_i_* represents for layer *i* the loss relative to the prediction error, *g_i_* is a sparse inducing prior that constrains the majority of the model neurons in *γ_i_* to be inactive and *a_i_* is a therm that scales the loss function and regulates the strength of the feedback. In this notation, we also have *γ*_0_ = *x* and *p*_0_ (*x*) = *x*. In order to solve the inverse problem, we use the *ℓ*_1_ norm (denoted ‖·‖_1_) instead of the *ℓ*_0_ pseudo-norm which is non-derivable. In a network composed of *N* hierarchical layers the total loss is given by:

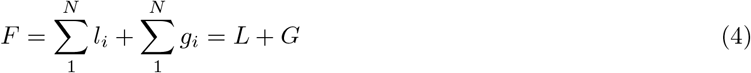

The best *γ_i_* for each layer is inferred through a multi-layer version of the Fast Iterative Thresholding Algorithm (FISTA) [32], introduced in [24]. This process is usually referred to as **inference**[33]. The update term is:

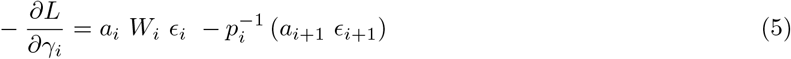

The left term in the above equation encodes for the reconstruction error, while the right one carries the feedback from the upper layer. 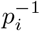 is defined as the unpooling function, that is, an approximation of the max pooling derivative; it is composed by a matrix filled with ones in the position of the local maximum elements and with zeros everywhere else. Finally, sparsity is obtained through a thresholding operator 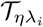. We also impose the elements in the neural maps *γ_i_* to be non-negative. The optimal state at the step *k* + 1 is given by:

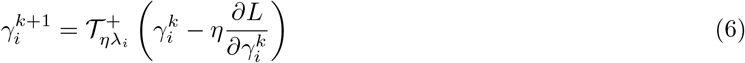

Where *η* is the time step of the inference process, *λ_i_* is the sparsity parameter and 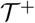 is the non-negative soft-thresholding operator (also known as a Rectified Linear Unit, or ReLU), defined as:

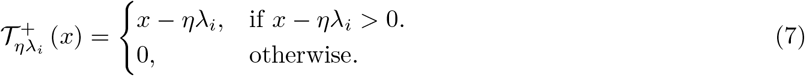

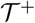 serves as both a rectified activation function and a sparsifying operator, forcing the majority of neurons in *γ_i_* to be inactive. Similarly, the synaptic weights can be updated by gradient descent:

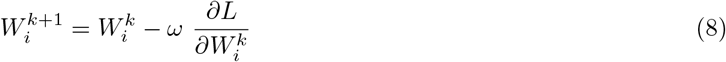

With *ω* being the learning rate for the weight update. It is worth noting that equation (8) represents a local Hebbian learning rule, as the update of *W_i_* only depends on the pre-synaptic (*γ_i_*_−1_) and post-synaptic (*γ_i_*) sparse activity:

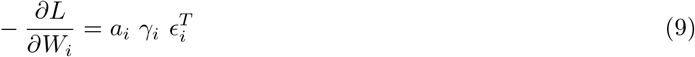

The optimization of *W_i_* is called **learning**. In this study, we model simple and complex-cells in a 2 layer SDPC network, with *γ_s_* encoding the activity of simple cells in the first layer and *γ_c_* for complex cells in the second layer. The loss function we wish to minimize is:

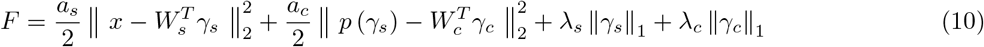

Fig. 1-a, shows the update scheme of the SDPC network used in this study.

### 2.2 Retinotopy and convolution

In the SDPC framework, we impose the weight matrix *W* to have a convolutional structure. Convolution is an efficient mathematical framework to model retinotopic activity in the visual system: synaptic weights are repeated across the image to tile the whole input image. This brings the advantage of having a model that is translation invariant: the synaptic weights (also called *features* or *kernels*) encoded by *W* can be freely translated across the input image. The resulting vector *γ* can thus be viewed as a *neural response map*, whose pixels encode for neurons sensitive to a specific feature and responding at a specific position in the input image, in perfect analogy with a retinotopic map (see Fig. 1, b). The convolutional structure is repeated for *M* different kernels, such as to define *M* different maps, called channels. Every channel represents the response of the identical neurons (sharing the same *kernel*) for each position in the input image. The convolution in each layer of the network is then defined by two terms:

- *Kernel size*: it indicates the size, in pixels, of the synaptic weights in *W*. Each pixel in *γ* can be mapped to a region equal to the kernel size in the input image.
- *Stride*: it is a measure of the amount of overlap of the convolutional kernels in the input image. Intuitively, for a given image *x*, the higher the stride the smaller will be the corresponding map *γ*.

Importantly, replacing classical matrix multiplication with convolution does not change the mathematical framework of SC and PC but rather makes the model scalable to different image sizes and imposes neural representations to be translation invariant. One limitation of convolution is that each plane of the *M* maps in *γ* is independent from the others. This makes convolution unable to account for interactions between different feature and to model higher order structures like orientation maps.

### 2.3 Inducing invariance by pooling

In section 2.1, we introduced the concept of a pooling function *p*(·) to model complex cells in the SDPC network, and we showed that the classical PC framework can be extended to take into account this kind of nonlinearity. Such a behaviour can be achieved with max-pooling, a function that selects the maximum response within a group of cells (here, *γ_s_*_1_ and *γ_s_*_1_ for an example):

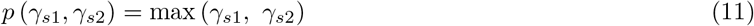

The max-pooling performs a winner-takes-all computation rather than calculating the energy of the firing of a pool of neurons [8]. We decided to use max-pooling as a way of generating invariance for two reasons: First, winner-take-all can account for numerous nonlinear responses in the sensory cortex, and it is regarded as one of the fundamental components of sensory processing in the brain [8, 12, 35]; Second, max-pooling is the most used form of spatial nonlinearity in convolutional neural networks [36] and this allows us to draw a parallel between classical feed-forward neural networks and deep predictive coding models like SDPC. Given a neural response map, *γ_s_*, max-pooling can be seen as a nonlinear convolution whose output, *p_s_* (*γ_s_*), is the maximum value in each pooling region (see Fig. 1, b). Just as convolution, max pooling can be defined in terms of kernel size and stride. In this study, we introduce three types of pooling function:

- MaxPool 2*D_S_*, acts in the spatial dimension, independently for each feature. In this study, we use a pooling kernel of 2 × 2 neurons with a stride of 2.
- MaxPool 1*D_F_*, selects the maximum across a neighborhood of adjacent planes of *γ_s_* along a 1*D* circular space. This function acts in the feature space, leaving the spatial encoding unchanged. For this function, we used a linear kernel of size 4 and stride equal to 1.
- MaxPool 2*D_F_*, arranges a feature space composed of *M* neurons on a grid of 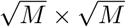 grid, for each spatial location. Then it selects the maximum activity across 2*D* pooling regions, with a size of 2 × 2 neurons with a stride of 1. This pooling function only acts in the feature space, just as the one we defined above.

Since the pooling functions operate in two different sub-spaces, we can apply them in sequence to combine the effect of the two strategies (see Fig. 1, b). We call these functions MaxPool 2*D_S_* + 1*D_F_* and MaxPool 2*D_S_* + 2*D_F_*.

### 2.4 Training

We built different SDPC networks to model simple and complex cells in V1. Each network had kernel sizes of 7 × 7 pixels for the first layer (*W_s_*), and 4 × 4 for the second layer (*W_c_*). For both layers, we used a stride of 1. We tested the network with 4 different pooling functions: MaxPool 2*D_S_*, MaxPool 2*D_F_* MaxPool 2*D_S_* + 1*D_F_* and MaxPool 2*D_S_* + 2*D_F_*. Importantly, we introduced a zero padding before the pooling to make sure that first and second layer cells have the same receptive field size with respect to the input image (14 × 14 pixels). In all the settings, *a_s_* and *a_c_* were set to 1 and 4 respectively. Each network was also trained with different neural populations sizes: with *M_s_* and *M_c_* equal to 36, 49, 64, 81, 100 and 121 neurons for the first and second layer respectively, leaving us with 36 networks for each tested pooling function. The parameters of the network *W_s_* and *W_c_* were initialized as random noise and were normalized during training such that the energy (Euclidean norm) of each kernel was set to 1. All the networks were trained using grayscale natural images from the STL-10 dataset (96 × 96 pixels per image) [37] for 9 epochs (28125 iterations with mini-batches of 32 images). The input images were pre-processed with a whitening filter similar to the one used in [16] to model retinal processing. Each image was then bounded between the values 1 and 1. The synaptic weight *W_s_* and *W_c_* were updated using stochastic gradient descent with a learning rate, *ω*, of 0.01 and a momentum, *β*, of 0.9. Additionally, the sparsity parameters *λ_s_* and *λ_c_* were gradually incremented during training up to a value of 0.1. The full model is implemented in PyTorch and available at https://github.com/XXX/YYY_ZZZ.

### 2.5 Modulation ratio and complex behaviour

In this study, in order to quantify the simple and complex behaviour of model neurons in the SDPC, we use the classical 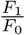 measure, also known as the modulation ratio [4]. Given the response of a neuron to a grating drifting at temporal frequency *f*, the 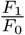 measure represents the ratio between the first harmonic of the response (*f* = *F*_1_) and the mean spiking rate *F*_0_. Intuitively, a high modulation ratio (between 1 and 2) indicates that the cell is sharply tuned to a specific phase and it is thus regarded as simple. If a cell shows a low modulation ratio (between 0 and 1) it is then regarded as complex, showing a broad tuning to the stimulus’ phase. For a neural population containing simple and complex cells, the distribution of the modulation ratios will appear to be bi-modal, with two peaks roughly in the regions described above. In [34] the authors showed that the modulation ratio can be derived analytically from a half-rectified model of the a neuron’s response to a drifting grating. In particular, 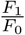 depends nonlinearly on *χ* defined as:

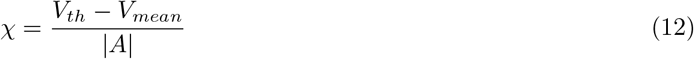

Where *V_mean_* and *V_th_* represent respectively the mean membrane potential and the threshold for spiking generation and *A* is the maximal amplitude of the modulation. Here we use the simplified rate based model:

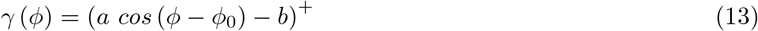

With *γ* being response of the model neuron, *φ* is the stimulus’ phase and *φ*_0_ the phase at which the response is maximal. Finally, ()^+^ indicates an half-rectified function that only output zero or positive values. In this case 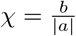 and the modulation ratio can be obtained through the nonlinear mapping (see [34] for details):

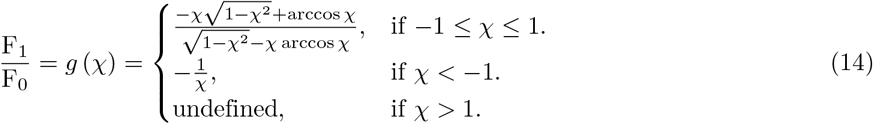

In [34], the authors suggest that the classification of V1 cells as simple or complex might be caused uniquely by the nonlinear relationship between 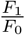 and *χ* rather than reflecting an actual neuro-physiological difference. Here, for simplicity, we refer to the same values conventionally used in literature: we consider model neurons as simple-like if 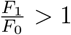 and as complex-like if 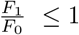. We use the modulation ratio also to evaluate the response to rotating stimuli in order to compare it to the response in the case of drifting phase. Although this measure is not in standard use to evaluate orientation tuning, it can be seen as a spectral analysis of the orientation response, similarly to the work of Wörgötter and Eysel [38]. Nevertheless, to perform statistical tests on the different conditions, we analyse the distribution of *χ* (see Supplementary information S.1.1). Indeed, *χ* is much easier to analyse as it does not show the typical bimodal distribution of 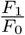 and is linked to 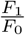 through a nonlinear, monotonous and invertible function (equation 14).

### 2.6 Local homogeneity index (LHI)

To evaluate the functional implication of the orientation maps learned by our model, we used the local homogeneity index (LHI) as defined by [39]. The LHI measures the similarity of the preferred orientation in neighboring cells:

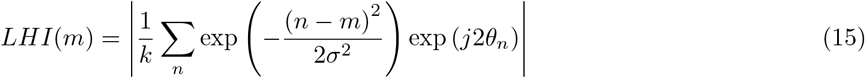

With *m* being a location on the orientation map, *n* a set of neighboring locations and *θ_m_* the preferred orientation at *n*. *j* is the imaginary unit, so that 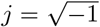 and represents the module of a complex number. k is constant that normalizes the measure, it can be change if *m* and *n* are bi-dimensional or mono-dimensional, for example in the case of MaxPool 2*D_F_* or MaxPool 1*D_F_*. Finally, *σ* is proportional to the width of the gaussian window in which the LHI is calculated, here we set *σ* = 1 pixel. The LHI is bounded between 0 and 1, with high values corresponding to iso-orientation domains and low values to pinwheels. We define pinwheels as local minima in local neighborhoods of 3 × 3 pixels with LHI under a threshold of 0.2. In order to calculate the pinwheel density we define the average size of a cortical column as the ratio 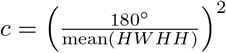, for each tested network. The pinwheel density is then defined as:

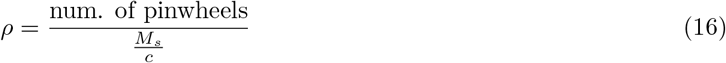

### 2.7 Log-Gabor fitting

To estimate the optimal frequency and orientation for each model neuron, we fitted the synaptic weights of each cell with log-Gabor wavelets [18, 40]. This fit allows us to identify the preferred stimuli for each neuron in the first and second layer. Specifically we can extract: the preferred orientation (*θ*), phase (*ϕ*), frequency (*f*_0_) as well as the the orientation tuning width (half width at half height, HWHH) [41]. Since second layer cells in the model project into the first layer of the network through a nonlinearity (pooling) we defined an approximated linear mapping 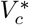 for the second layer such that:

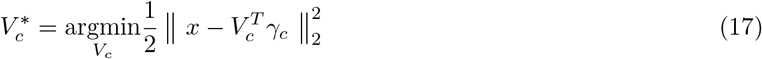

Specifically, 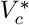 is used to back project (and visualize) the second layer’s weights in the input space. The kernels in 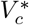 were then fitted using log-Gabor wavelets to assess for the best orientation and spatial frequency to stimulate the second layer of the network.

### 2.8 Drifting grating vs. Rotating grating

One goal of our study is to assess the ability of our network to predict complex behavior in V1, that is, to show phase-invariant responses. We test the network’s invariance to drifting and rotating gratings. We do this to make sure that the networks we model are show indeed responses invariant to phase but that they remain tuned to orientation, as observed in neurophysiological experiments. Using the preferred orientation, *θ*, and frequency, *f*_0_, for each neuron in the first and second layer, we created sinusoidal grating at the right bandwidth for each neuron. All stimuli were masked by a circular window of 14 pixels in diameter, that is the size of the receptive field of model neurons. Finally we evaluated the response of the network at different phases and orientation of the same grating (see Fig. 4).

## 3 Results

### 3.1 Pooling in a predictive coding network

The first important result of this study is the convergence of the SDPC algorithm in all the tested network settings: the SDPC efficiently learned translation-invariant representations for the naturalistic stimuli used during training: in all tested conditions, the network reached a stable point in terms of the loss function value (see eq. 4). This was the case for the different network sizes (36, 49, 64, 81, 100 and 121 neurons for each layer) and the different combinations of pooling functions: MaxPool 2*D_S_*, MaxPool 2*D_F_*, MaxPool 2*D_S_* + 1*D_F_*, and MaxPool 2*D_S_* + 2*D_F_*. As in [23] and [24], the network efficiently developed edge-like receptive fields (RFs) in the first layer (see Fig. 1,b). Inferring the shape of the second RFs is less trivial, indeed the pooling function *p_s_* (*γ_s_*) makes it challenging to project the learned filters in the space of the input image (see [23] for details). In Fig. S1 we show *V_c_*, a linear approximation for the second layer kernels (see Methods 2.7). From these linear projections, we can see that the RFs in *W_c_* also have the shape of localized edge-like filters (see Fig. S1). Nevertheless, as we introduce the nonlinear function *p_s_* (*γ_s_*), the response of the neural map *γ_s_* will be strongly nonlinear and show different degrees of robustness to changes in the input stimuli. We will analyze these results in detail in the following sections.

### 3.2 Phase invariance and complex behavior

Complex cells are present in all networks with *p_s_*=MaxPool 2*D_S_*, *p_s_*=MaxPool 2*D_S_* +2*D_F_* and *p_s_*=MaxPool 2*D_S_* + 1*D_F_*. Indeed, in the second layer neurons are invariant to changes in the stimulus’ phase but still selective to orientation (see Fig. 2). To quantify phase invariance, we use the modulation ratio 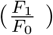 introduced by [4] (see Methods 2.5). A cell is classified as simple if 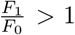 and as complex if 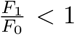. In these three settings, the distribution of 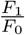 shows that the first layer develops exclusively simple-like neurons, while the second layer shows a broader distribution with the presence of complex-like neurons depending on the pooling function used (Fig. 3, fourth column). When spatial pooling and feature pooling are combined together (for *p_s_* =MaxPool 2*D_S_* + 1*D_F_* and *p_s_* =MaxPool 2*D_S_* + 2*D_F_*), the second layer of the network develops a large number of complex-like neurons, which explains the higher peaks at 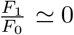 (see Fig. 3, c-d). Interestingly, these two conditions correspond to networks that develop a topographic map in the orientation space, but not in the phase space. Indeed, we observe that for *p_s_* =MaxPool 2*D_F_*, our model does not develop complex cells, i.e. the 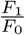 is high for both layers. Similar distributions of complex neurons are observed for *p_s_* =MaxPool 2*D_S_* but, in this condition, the network does not develop a cortical map.

**Fig 2.**
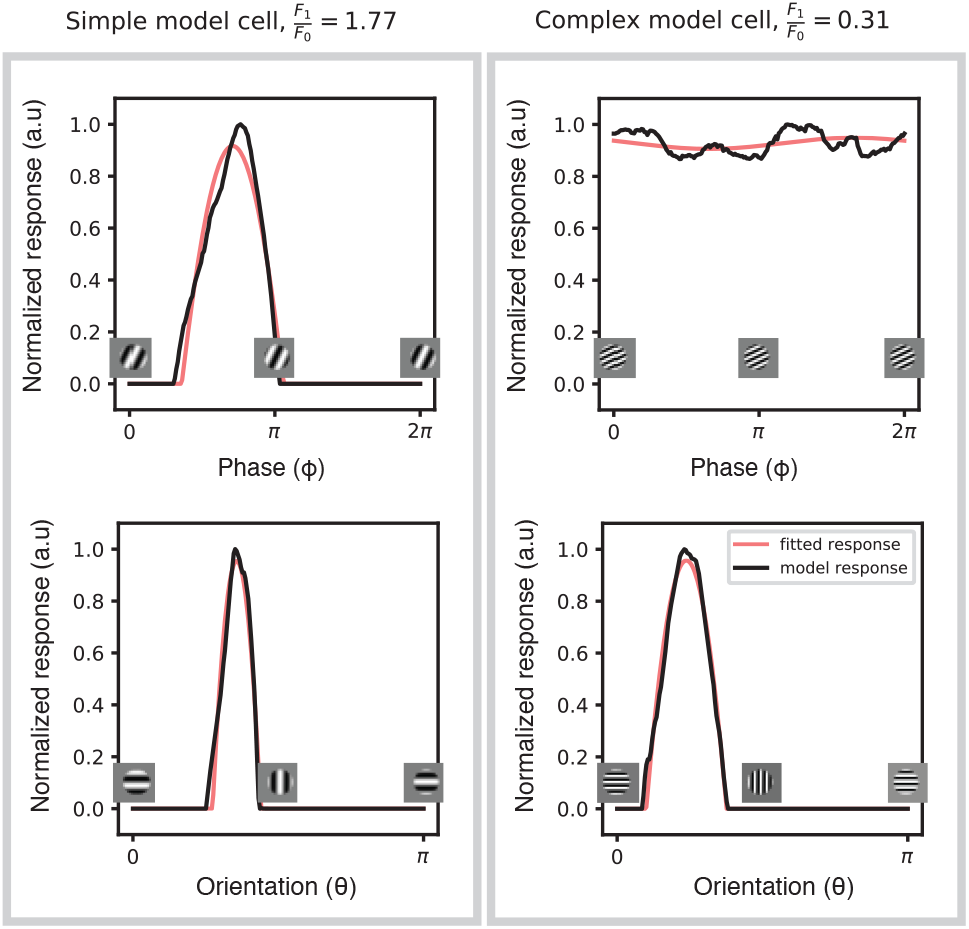
Response of simple-like and complex-like model neurons. Example of two model neurons exhibiting a simple (**left**) and complex (**right**) behavior. The black lines indicate the model neuron’s response when its receptive field contains a drifting (**top**) or rotating (**bottom**) stimulus; the red lines indicate the response modeled according to a half-rectified sinusoidal model as in [34]. The modulation ratio, 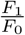, is greater than 1 for simple neurons that are tuned to a specific phase. A complex response, that is partially or completely independent to phase, is quantified by a low modulation ratio (less than 1). Importantly, both neurons remain tuned to orientation.

**Fig 3.**
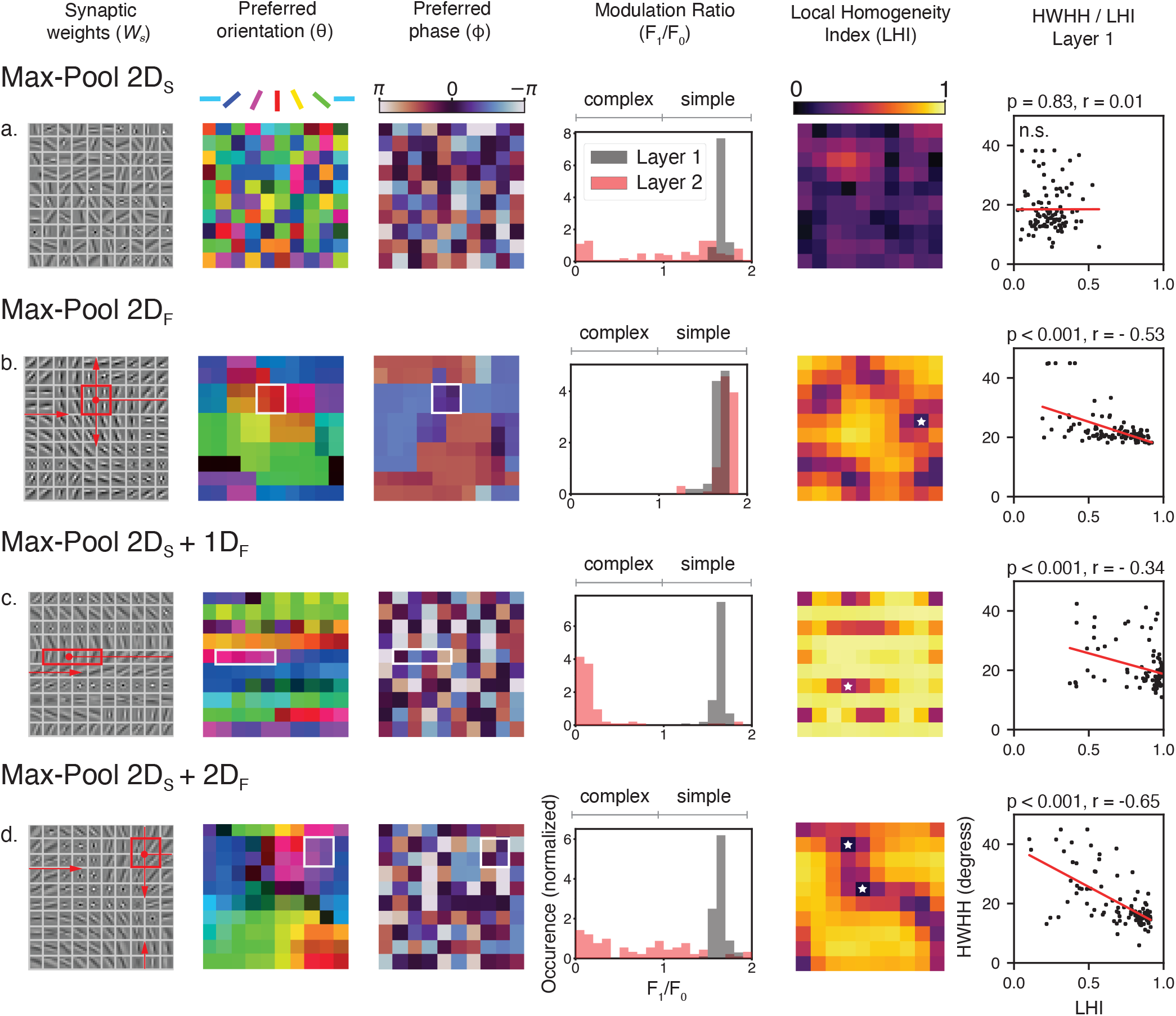
Emergence of orientation maps in the first layer and complex response in the second layer. We show the different properties learned by our model depending on which pooling functions we used (**a**-**d**). All networks showed here are trained with *M_s_* = 100 and *M_c_* = 100. **First column**. Representation of the synaptic weights of the first layers *W_s_*. The red rectangles for (**b**-**d**) indicate the size of the feature pooling, the red arrows illustrate the circular (**c**) or toroidal (**b** and **d**) structure of the feature space. **Second column**. Orientation preference of each neuron represented by the filters in *W_s_*. The luminance of the map is modulated by the orientation selectivity (HWHH) of each neuron. **Third column**. Phase preference of each neuron represented by the filters in *W_s_*. **Fourth column**. Distribution of the modulation ratio 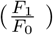 for the first and second layer of the network. **Fifth column**. Local homogeneity index (LHI) associated with each element is *W_s_*. White stars indicate the position of pinwheels. **Sixth column**. Linear relationship between the LHI and the orientation selectivity, measured by the half width at half height (HWHH) of the response of cells from the first layer.

### 3.3 Learning topographic orientation maps in an SDPC network

Networks trained using pooling functions that act in the feature space (MaxPool 2*D_F_*, MaxPool 2*D_S_* + 2*D_F_*, and MaxPool 2*D_S_* + 1*D_F_*), learned different topographic structures for *W_s_* and, by consequence, *γ_s_*. Interestingly, as we introduce pooling in the feature space (see Methods 2.3), the arrangement of filters on *W_s_* converges to form a topographic orientation map where neighboring neurons become tuned to similar orientations. To compare quantitatively the orientation maps learned by our model with the ones observed in neurophysiological experiments, we used the local homogeneity index (LHI), introduced by [39] (see Fig. 3, b-c-d, second column). We observed that the orientation maps developed by the SDPC contain regions where orientation preference varies smoothly (high LHI) combined with local discontinuities (low LHI), in analogy with pinwheels observed in higher mammals [27–29].

One may wonder if the existence of these topographic maps has functional implications in the properties of the first layer neurons. Indeed, some electrophysiological experiments have shown a clear link between a neuron’s position in the cortical map and its tuning properties [39, 42] (but see [28]). Specifically, neurons located near pinwheels tend to have a broad orientation tuning, while neurons located in iso-orientation regions display a narrower, more selective tuning. We evaluated the relationship between the LHI of single neurons in the first layer and their orientation tuning, evaluated as the half-width at half-height (HWHH) of their response to a rotating grating. We found the linear relationship already found in [39] and [42]. This relationship is valid for all networks with a 2*D* topographic map (*p_s_* =MaxPool 2*D_S_* + 2*D_F_*) and *M_s_* ≥ 64 (*p* < 0.01). The linear relationship is significant (*p* < 0.01) for few configurations of *p_s_* =MaxPool 2*D_F_* and *p_s_* =MaxPool 2*D_S_* + 1*D_F_*, with no clear dependence on *M_s_*.

Interestingly, the networks trained with a combination of spatial and feature pooling (MaxPool 2*D_S_* + 1*D_F_* and MaxPool 2*D_S_* + 2*D_F_*) appear to develop a topographic map when we look at the preferred orientations *θ* of neurons in the first layer (Fig. 3, second column). However, this structure is not present if we look at the preferred phase map *ϕ*, where there appear to be no particular structure (Fig. 3, third column), in line with the known neurophysiology [43]. This is not true in the case of MaxPool 2*D_F_*, where both maps appear to be organized in groups of cells sensitive to similar orientations and phases, in contrast with neurophysiological evidence. In the following sections, we will analyze the implication of these results.

We used the LHI to evaluate the number of pinwheels generated by our model in different tested conditions (see Methods 2.6). We found that the tested pooling functions, MaxPool 2*D_F_*, MaxPool 2*D_S_* + 2*D_F_*, and MaxPool 2*D_S_* + 1*D_F_*, generated a comparable number of pinwheels centers even for networks about 4 times bigger that the smallest network tested: *M_s_* = 36 and *M_s_* = 121 (see Fig. S2). Thus, the number of pinwheels singularities remarkably does not depend on the size of the first layer *M_s_*, with a value in the same order of magnitude of *π* [44].

### 3.4 Invariance to phase vs. invariance to orientation

We want to explore further how is it that the complex cells developed by the second layer of our model show invariance to the stimulus phase while remaining tuned to orientation (Fig. 4). A highly nonlinear network could respond unspecifically to a broad set of different stimuli. Invariance to multiple properties of the stimuli (e.g. both to phase and orientation) would be in contradiction with neurophysiological experiments. In fact, it is well established that complex cells exhibit phase invariance but show orientation tuning similar to simple cells [4, 5]. To assess whether the network’s response is orientation-selective or not, we tested the network’s invariance property to both phase and orientation of gratings. We used drifting gratings as stimuli that change phase over time. Conversely, we used rotating gratings that change orientation, to modulate the response of orientation-selective neurons. This was performed for each cell of the second layer of our network at its optimal spatial frequency (see Methods 2.8). For quantification purposes, we define R as the percentage of cells that show a complex-like response for phase, or that are unselective to orientation 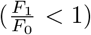. We report the results relative to the second layer of the networks, as in all the tested conditions, the first layer of the network exclusively exhibited simple-like responses (*R_ϕ_* = 0, see Fig. 3, fourth column). In Fig. 4, the black and green lines *R_ϕ_* and *R_θ_* correspond to the dependence on phase (drifting) and orientation (rotating), respectively. We analyze these results as a function of the number of channels in the first layer, *M_s_* to evaluate the impact of the network structure on the second layer’s responses. For *p_s_* =MaxPool 2*D_S_*, the networks showed high *R_ϕ_* (up to 60%) and low *R_θ_* (at most 20%), independently to *M_s_*. This suggests that the SDPC network efficiently develops phase-invariant responses while maintaining a relatively narrow tuning to the stimulus’s orientation (see Fig. 4, top-right). When *p_s_* =MaxPool 2*D_S_* + 1*D_F_*, that is, a spatial pooling and a feature pooling organized on a linear structure, *R_ϕ_* is much higher (up to 100%, see Fig. 4, bottom-right) and a smaller *R_θ_*, also independent on *M_s_*; this can be explained by the combined action of pooling across spatial locations and across features with same orientation but different phases (see Discussion and Fig. 3). Interestingly, only when we impose *p_s_* =MaxPool 2*D_S_* + 2*D_F_*, the network’s behavior appears to vary as a function of *M_s_* (see Fig. 4, bottom-left). For low dimensions (up to *M_s_* = 64), the tested networks show a high fraction of complex-like cells. For *M_s_ >* 64, *R_ϕ_* decreases as *M_s_* increases (see Supplementary Info. S.1.1 for statistical tests). The same effect can be observed for *R_θ_*, which appears to be high for *M_s_* = 36, and then decreases with increasing *M_s_*. Overall, none of the observed behaviors did depend on number of neurons in the second layer (*M_c_*), for any of the tested conditions (not shown).

**Fig 4.**
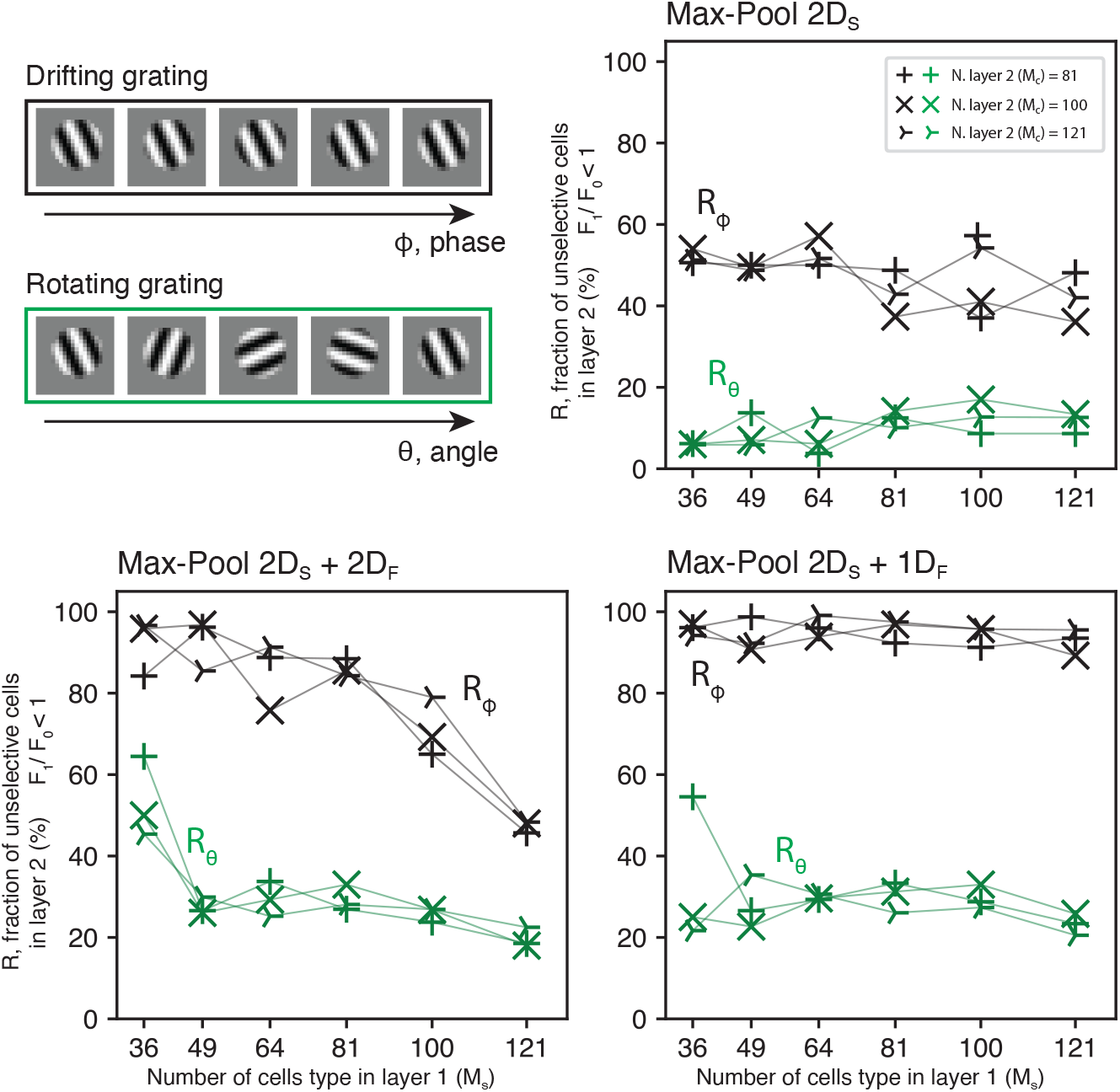
Complex cells population analysis. Here we show how the type of pooling function used and the number of cells type present in the first and second layers of the network (*M_s_* and *M_c_*, respectively) affect the complex cells population’s properties developed by the model. **Top-left.** We tested the network’s second layer invariance to phase (drifting) and orientation (rotation) in sinusoidal gratings (see Methods 2.8). The ratios of cells *R_ϕ_* (complex) and *R_θ_* (orientation invariant), are defined as the percentage for which 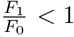 in the second layer of the network. While the network’s second layer shows a strong invariance to a drifting grating (black lines), the same effect is not present for the rotating grating (green lines). This result indicates that the network’s invariant response is specific to the stimulus’s phase and not to its orientation. Overall *R_ϕ_* is much higher when the max-pooling acts also in the feature space (*p_s_* =MaxPool 2*D_S_* + 1*D_F_* and *p_s_* =MaxPool 2*D_S_* + 2*D_F_*). Interestingly, for *p_s_* =MaxPool 2*D_S_* + 2*D_F_*, the number of complex-like cells seems to depend on the size of the network’s first layer *M_s_*.

## 4 Discussion

In [23], we introduced the SDPC algorithm, and we used this framework to model local interactions in the early visual cortex (V1/V2). We showed the SDPC can predict how strong intra-cortical feedback connectivity (from V2 to V1) shapes the response of neurons in V1 according to the Gestalt principle of good continuation. Here, we show that the SDPC algorithm can be extended to account for highly nonlinear operations in cortical processing within V1. Our goal in this study was to implement a 2-layer model of V1 to explain the emergence of two distinct populations of neurons: simple and complex cells. In particular, we introduced a max-pooling operation between the network’s layers to increase the generalization capability of our model (see Fig. 1, a). Thanks to this nonlinear stage, the SDPC model can account for nonlinear neural responses, in analogy to complex cells in V1. We tested and combined the effect of three classes of max-pooling functions: MaxPool 2*D_S_*, MaxPool 1*D_F_* and MaxPool 2*D_F_*; the first allowed our model to pool responses across spatial locations, the other two to pool across feature space.

### 4.1 Results summary

Here we summarize the results obtained by training the SDPC model with four different sets of pooling functions, for the first and second layer of the neural network: For MaxPool 2*D_S_*, the network only pools in the retinotopic space, and no orientation map is learned in the first layer. Nevertheless, the model efficiently develops complex cells, as observed in rodents, in its second layer (see Fig. 3, a and Fig. 4). This network structure can be seen as a model of rodents’ visual cortex, as they lack orientation maps. In the case of MaxPool 2*D_F_*, the network only pools in the feature space. In this condition, the model learns a topographic map in the first layer for both orientation and phase (Fig. 3, b). This condition has no physiological basis and is only used as a control condition. Indeed, in this configuration, no complex cells are present in the second layer for any tested condition (*R_ϕ_* ≈ 0%), and cortical maps for phase have never been described. The two last pooling conditions can be seen as an equivalent of carnivores’ or primates’ visual cortex. For MaxPool 2*D_S_* + 1*D_F_*, the network pools in both retinotopic and feature space (circular topology). In this case, the first layer of the network develops a topographic structure for orientation preference but not for phase (Fig. 3, c). The second layer presents a high fraction of complex cells (up to *R_ϕ_* = 100%) in all tested conditions irrespective of *M_s_*. For MaxPool 2*D_S_* + 2*D_F_*, the network pools in both retinotopic and feature space (toroidal topology). Similar to the previous case, the network develops, in the first layer, a topographic map for orientation but not for phase (Fig. 3, d). In this case, however, *R_ϕ_* appears to depend on *M_s_*, with smaller first layers (low *M_s_*) appearing to produce more complex cells (Fig. 4). Thus, a simple pooling rule can explain the emergence of both complex cells and cortical maps in higher mammals.

### 4.2 Relation to the state-of-the-art

Our work proposes a novel framework to understand the emergence of complex cells’ properties through pooling in a topographically organized network. It builds upon existing literature but differs in some essential points. First, Karklin and Lewicki [45] studied the emergence of complex cells using a probabilistic framework. However, their model is equivalent to a single-layer network, thus not addressing the hierarchical nature of phase invariance. Hyvarinen *et al.* [30] defined a network that is specifically designed for a 2-layered structure. This sparse coding model explicitly minimizes the local pooled energies of groups of complex cells. In contrast, in the SDPC the topographical organization is not imposed but emerges as a consequence of the prediction error generated by the second layer. Similarly, Hyvarinen *et al.* [31] defined a probabilistic framework to model the residual statistical dependencies in the latent variables of an Independent Component Analysis (ICA) framework. This latter model bears some similarities with ours. But crucially, rather than explicitly modeling the statistical relationships in a one-layer network, we separate the generative problem into multiple sub-functions, each belonging to a specific layer, according to the predictive coding principle. This process can be summarized in two steps. First, we allow the transfer of information between layers thanks to the forward and feedback connections. In this stage, the hidden relationships in the latent variables (neurons’ firing) can be inferred thanks to the feature pooling function we defined in section 2.3. Second, these dependencies are predicted by the second layer’s activity, whose variables are regarded as independent. This structure allows the model to grasp higher-order relationships between the variables while leaving the core PC framework unchanged and making it extendable to an arbitrary number of layers. The networks defined in [30] and [31] are composed of neurons with relatively large receptive fields (32 × 32 pixels). Most importantly, these models are based on a fully connected architecture, which means that each neuron has learnable synaptic weights that span the whole size of the input image. This type of architecture does not allow to disentangle the role of retinotopy from the one of orientation map in building complex cell responses, as the two structures are merged in the same map. As a consequence, the models listed above do not explain the emergence of complex cells in animals that do not show orientation maps, such as rodents [27, 29]. On the other hand, our model is based on a convolutional structure that makes the algorithm translation invariant and scalable to any input size (see Methods 2.2). The convolutional structure of the SDPC allows evaluating separately retinotopy (convolution and spatial pooling) and orientation maps (feature pooling). Lastly, we would like to emphasize that another major novelty of our proposed model is interpreting complex cells’ properties within the predictive coding framework.

Similar to [14] and [15], our network is based on multiple layers encoding the hierarchical structure of the input data. In these studies, the authors define a multi-layer sparse network to model contour-sensitive cells in the early visual cortex (V2). Hierarchical sparse coding and the SDPC share the same generative problem (see [24] for details). However, the SDPC can account for feedback connections that carry information backward in the network during inference. In previous works, we showed that a feedback information stream offers significant advantages to classical multi-layer sparse coding networks. These effects are relevant both in terms of efficiency [24] and plausibility of the neuroscientific modeling [23]. Besides, the models defined in [14] and [15] are based on an energy model in the Fourier space while, in the SDPC, we learn the synaptic weights of the network directly from the input data. For a detailed discussion of the critical points of our model see Supplementary information S.1.2.

### 4.3 Contributions

The pooling stage, *p_s_*(*γ_s_*), allows the SDPC model to create representations that are invariant to local changes in the spatial (retinotopic) domain (MaxPool 2*D_S_*) and in the feature space (MaxPool 1*D_F_* and MaxPool 2*D_F_*). The prediction generated by the second layer 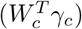 inherits the same invariant structure of *p_s_* (*γ_s_*), allowing the network to learn representations that maximize invariance by minimizing the prediction error *E_c_* (see section 2.1). We found that our model of V1 learns RFs in the form of localized, band-pass, orientation-selective filters. This result is an important sanity check for any unsupervised sparse network used to model the behavior of V1. Indeed, sparse networks are known to efficiently predict the shape of V1 RFs in higher mammals [16, 18]. The SDPC network generates cells tuned to specific orientation and spatial frequency in both the first and second layers. We can state that the SDPC framework can be efficiently extended to include highly nonlinear functions (max-pooling) while making valid predictions on the structure of V1 receptive fields and the shape of V1 neural responses.

#### 4.3.1 SDPC predicts the emergence of cortical orientation maps

We observed that when the pooling function used in the model acts in the feature space (MaxPool 2*D_F_*, MaxPool 2*D_S_* + 1*D_F_*, and MaxPool 2*D_S_* + 2*D_F_*), the SDPC model develops a topographical orientation map for the RFs of the first layer. These orientation maps show quantitatively strong similarities with those observed in higher mammals: First, we found that a linear relationship exists between the local homogeneity index (LHI) of a model neuron (i.e. its position relative to pinwheels in the map) and its orientation tuning as observed in neurophysiological studies [39, 42]. In particular, neurons near pinwheels (low LHI) have a broader tuning than those in iso-oriented regions (see Results 3.3). Second, we found the number of pinwheels for each of the conditions listed above to be independent of the network first layer’s size, *M_s_* (Fig. S2). This result is in line with previous neurophysiological studies that found pinwheel density to be constant across species (circa 3.14 singularities per cortical column), regardless of the size of V1 [27, 44] (but see [46] for a detailed computational model). Our model did not converge to the same value, but the same order of magnitude, as in our study orientation maps and cortical maps are generated by two different pooling functions. Nevertheless, we found that the density of pinwheels remains constant even in networks four times bigger than the smallest one we tested. To sum up, our model efficiently predicts a key property of cortical orientation maps across different species, carnivores, and primates, that is the emergence of pinwheels and orientation domains with a constant pinwheel density even for a large diversity of V1 sizes [27, 44, 46]. The most remarkable fact is that, in our model, this structure is not enforced but learned directly from the data in an unsupervised manner (see Fig. S3), and it only depends on the type of pooling function used. But how could this particular topography be learned? The response of the model is not driven by a feed-forward transform but by a prediction mechanism. In other words, the second layer of the network learns representations that are invariant thanks to the max-pooling stage. The elicited prediction error, in turn, forces the representation in the first layer to align with the top-down prediction. In short, the predictive coding mechanism induces a topographic organization through feedback connectivity. Moreover, the SDPC solves a convex optimization problem and is, thus, guaranteed to converge to a near-optimal solution [32]. This indicates that *orientation maps are the most cost-efficient representation, under the predictive coding principle*. Thus, this simple computational framework offers an elegant explanation for the existence of orientation maps in higher mammals, as observed in electrophysiological experiments [5, 28], as well as a computational principle that explain their emergence [47].

#### 4.3.2 SDPC explains the role of cortical maps across species

A second important result resides in the fact that SDPC predicts the emergence of complex cells as a result of hierarchical processing. In fact, no complex-like cells were present in the first layer of the network in any tested conditions (Fig. 3, fourth column). This is in line with neurophysiological studies on gray squirrel [48], rabbit [49], cat [50] and macaque monkey [2, 18, 51], that found that simple cells are more common in the input layer of V1 (named layer 4), corresponding to the first layer of our network. Conversely, complex cells appear to be more common in superficial layers (named layers 2/3), corresponding to the second layer of our network, suggesting the existence of a hierarchical mechanism (see [52] for a detailed review). However, complex cells can also be found in the early stage of the primary visual cortex and have been shown to receive direct thalamic input [52], our model fails to predict the emergence of this specific population. We can argue that the responses predicted by our model can grasp some critical aspects of the neural processing in V1, like nonlinearities and hierarchical processing, but are still subject to simplifications due to computational constraints. We were able to efficiently model at least one component of the complex cells’ population in V1: layer 2/3 receiving input from layer 4. According to our model, complex cells emerge by pooling in the retinotopic space, even in absence of orientation maps, thanks to hierarchical pooling in V1. The same pooling principle could be applied directly to thalamic input generating first-order complex cells, as observed in the rabbit (see [53]). Indeed, we observed complex responses in networks that are at least able to pool in the retinotopic domain (MaxPool 2*D_S_*); this is in line with our hypothesis on the emergence of a class of complex cells as the result of pooling simple cells responses across a retinotopic organization (see Fig. 1, b) and with the fact that rodents and lagomorph show no orientation maps despite a huge diversity in V1 sizes [27, 29].

Nevertheless, the network can learn more efficient strategies to build up phase invariance by pooling across the feature space. Indeed, when *p_s_* =MaxPool 2*D_S_* + 1*D_F_* (see Fig. 3, c), the network develops an orientation map where the preferred orientation of adjacent neurons varies gradually along a circular structure (ring). Interestingly, this type of structure does not appear in the phase space, where there is no particular organization. The same computational principle can be extended to a 2*D* topology by using *p_s_* =MaxPool 2*D_S_* + 2*D_F_*. There, the network can exploit an additional degree of freedom in the feature space and learns topographic orientation maps that show strong similarities to those observed in the primary visual cortex of higher mammals, as we discussed earlier. In this case, too, there is no organization in the phase domain. This lack of structure implies that for each pooling region in the feature space (red rectangles in Fig. 3, b-d) the network pools over the responses to oriented features with different phases, generating phase invariance. An analogous mechanism was suggested to explain binocular disparity in cat’s complex cells [54, 55]. Interestingly, networks trained with only feature pooling (MaxPool 2*D_F_*) were unable to explain the emergence of complex cells, although this pooling mechanism generates orientation maps. We can observe that the features produce a topographic organization in both orientation and phase space, in contrast with neurophysiology(see Fig. 3, b). This lack of complex response is in line with our previous statement, as this topographic map does not allow the network to pool neural response in the phase space efficiently. This result is not surprising as, in the case of MaxPool 2*D_F_*, the networks do not perform pooling in the retinotopic space. This condition is not physiologically plausible as retinotopic maps are a structure common across all mammals [27, 29, 56]. Pooling across retinotopic positions (MaxPool 2*D_S_*) can explain, by itself, the emergence of complex cells by **position invariance**, a fundamental mechanism observed in complex cells [54]. In this case, the absence of cortical orientation maps suggests that the emergence of complex cells depends solely on pooling across different positions in the retinotopic space. This type of network could be the one preferably implemented in animals that do not exhibit orientation maps [27, 29]. This is in line with the results from [57], showing that orientation selectivity can emerge even in networks that connect locally neurons with a random preferred orientation, as in a salt-and-pepper map; in analogy with the random maps that the network produces in the case of MaxPool 2*D_S_* [57, 58]. On the other hand, in the case of MaxPool 2*D_S_* + 1*D_F_* and MaxPool 2*D_S_* + 2*D_F_* our model can generate **feature invariance** by pooling in the feature space. By consequence, the model converges to a configuration where complex cells can be generated by pooling across the same orientations and different phases (Fig. 3, c, d). In fact, in the Result sections 3.2 and 3.4, we showed that networks that develop topographic maps, on top of classical spatial pooling, develop more complex-like cells and, in general, more phase invariant responses, see in particular Fig. S4 and Supplementary Information S.1.1 for a detailed analysis. These results allow us to expand our previous statement: *orientation maps represent the most cost-efficient way to generate nonlinear complex cell responses, under the predictive coding principle*. These findings suggest that rodents should show a lower fraction of complex cells in V1, compared with other mammals. This prediction is, indeed, in line with neurophysiological findings in mouse [56, 59] and the rabbit [49], although squirrels (that are highly visual rodents) show a fraction of complex cells comparable to other mammals [58]. The most remarkable fact is that these results are obtained solely by changing a single computation mechanism, that is the type of nonlinear pooling used, and by learning neural responses directly from natural images. Altogether, these results suggest that SDPC could be regarded as a unifying mechanism in mammals’ V1, explaining the emergence of complex cells in all types of topographical structure (salt-and-pepper and orientation maps) as described in different species.

#### 4.3.3 Non linear representations depend on structure

Lastly, in section 3.4 we demonstrated that, although our model contains multiple sources of nonlinearity, it efficiently accounts for complex cells that are insensitive to the stimulus’ phase but remain significantly tuned for orientation. This result is significant for two reasons: first, this is in line with those found in electrophysiological studies [4, 5]. Second, it confirms that the network does not generate unspecific responses due to a high degree of nonlinearity. We observed another interesting phenomenon: in the case of a 2*D* orientation map (*p_s_*=MaxPool 2*D_S_* + 2*D_F_*) the number of complex cells developed by the model (as quantified by *R_ϕ_* in Fig. 4) appears to be inversely proportional to the number of cell in the first layer of the network (*M_s_*). This effect can be interpreted as: Networks with a limited number of neurons in the first layer are constrained to learn more nonlinear representations. At the same time, as the local neural population increases, the network is allowed to generate more diverse cell types, thus converting to more linear, simple-like responses. This phenomenon could explain different neural behaviors across species and cortical locations. Although the presence of topographic maps cannot be predicted by the size of the visual cortex [27], our results here suggest that salt-and-pepper organization could be linked to a particular pooling strategy. An interesting result in this sense was obtained by [60]: the authors were able to model the emergence of topographic maps by modulating the density of cell population between the lateral geniculate nucleus (LGN) and V1 (see also [53]): This simple model explains the different columnar organizations across species, from salt-and-pepper in rodents to orientation maps and pinwheels in primates. A similar principle could be translated to the SDPC model presented in this paper but in relation to the connectivity between layer 4 and layers 2/3 of V1. A possible hypothesis is that cortical regions show different types of topographic organization depending on the number of neurons they receive input from. For example, rodents could use unspecific connectivity (large *M_s_*) to generate a highly nonlinear response without a topographic map. At the same time, higher mammals could exploit, under some evolutionary wiring constraints, more adapted connections (lower *M_s_*) to fully exploit the presence of pinwheels and iso-oriented domains to generate phase-invariant responses within layer 2/3 complex cells. This mechanism could give an elegant and simple explanation for the different structural and functional aspects of V1 across species; however, such a hypothesis remains to be validated.

## S.1 Supplementary Information

### S.1.1 *Analysis on* χ

We define *χ_θ_* and *χ_ϕ_* as the measures obtained from the responses to rotating (angle, *θ*) or drifting (phase, *ϕ*) gratings respectively (green and black lines in Fig. S4). We use non parametric tests as both *χ_θ_* and *χ_ϕ_* do not follow a Gaussian distribution. We found that *χ_θ_* was significantly higher (less complex) than *χ_ϕ_* (one-tailed Wilcoxon signed-rank test) with *p* < 0.001 in all tested conditions. The only exception being for *p_s_* =MaxPool 2*D_F_* (not shown), as in this case, no network shows cells with 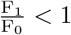 in the second layer. It is worth noticing that the median *χ_ϕ_* tends to be much lower when the pooling function includes the feature space, for *p_s_* =MaxPool 2*D_F_* and *p_s_* =MaxPool 2*D_S_* +1*D_F_*, than for a simple spatial pooling, *p_s_* =MaxPool 2*D_S_* +2*D_F_* (see Fig. S4). To assess the significance of *χ_ϕ_* and *χ_θ_* generating more complex-like or simple-like behavior we used a one-tailed binomial test (sign test). We set the null hypothesis *median*(*χ_ϕ_*) < −1 and *median*(*χ_θ_*) ≥ −1. Indeed, for *p_s_* =MaxPool 2*D_S_*, the median of *χ_ϕ_* is found to be significantly lower than *−*1 in only a couple of conditions, whereas in general it tends to be greater than *−*1. In analogy with the results shown above, for *p_s_* =MaxPool 2*D_S_* + 1*D_F_* the network significantly generates a majority of complex-like cells *χ_ϕ_* < −1. In the case of *p_s_* =MaxPool 2*D_S_* + 2*D_F_*, we found again a dependence of the amount of complex-like cells and *M_s_* with *χ_ϕ_* being significantly less than −1 for *M_s_* ≤ 100 and highly significantly (*p* < 0.001) for *M_s_* ≤ 81. On the other hand, *χ_θ_* is found to be significantly greater than −1 for all networks with *p_s_* =MaxPool 2*D_S_* + 2*D_F_*, *p_s_* =MaxPool 2*D_S_* + 1*D_F_* and *M_s_ >* 49. This indicates that, for *p_s_* =MaxPool 2*D_S_* + 2*D_F_* and *p_s_* =MaxPool 2*D_S_* + 1*D_F_* low *M_s_* tends to create highly nonlinear networks with broad tuning to both phase and orientation, while increasing further *M_s_* generates networks with more simple-like behaviors (see Discussion). For *p_s_* =MaxPool 2*D_S_* and *p_s_* =MaxPool 2*D_F_*, *χ_θ_* is significantly greater than −1 in all tested conditions. Finally, we found *χ_ϕ_* to be significantly smaller for *p_s_* =MaxPool 2*D_S_* + 1*D_F_* than *p_s_* =MaxPool 2*D_S_* in all tested conditions (single-tailed Mann-Whitney U test, *p* < 0.05). Whereas *χ_ϕ_* was significantly smaller for *p_s_* =MaxPool 2*D_S_* + 2*D_F_* than *p_s_* =MaxPool 2*D_S_* for all *M_s_* ≤ 100 and highly significantly (*p* < 0.001) for all *M_s_* ≤ 64.

**Fig S1.**
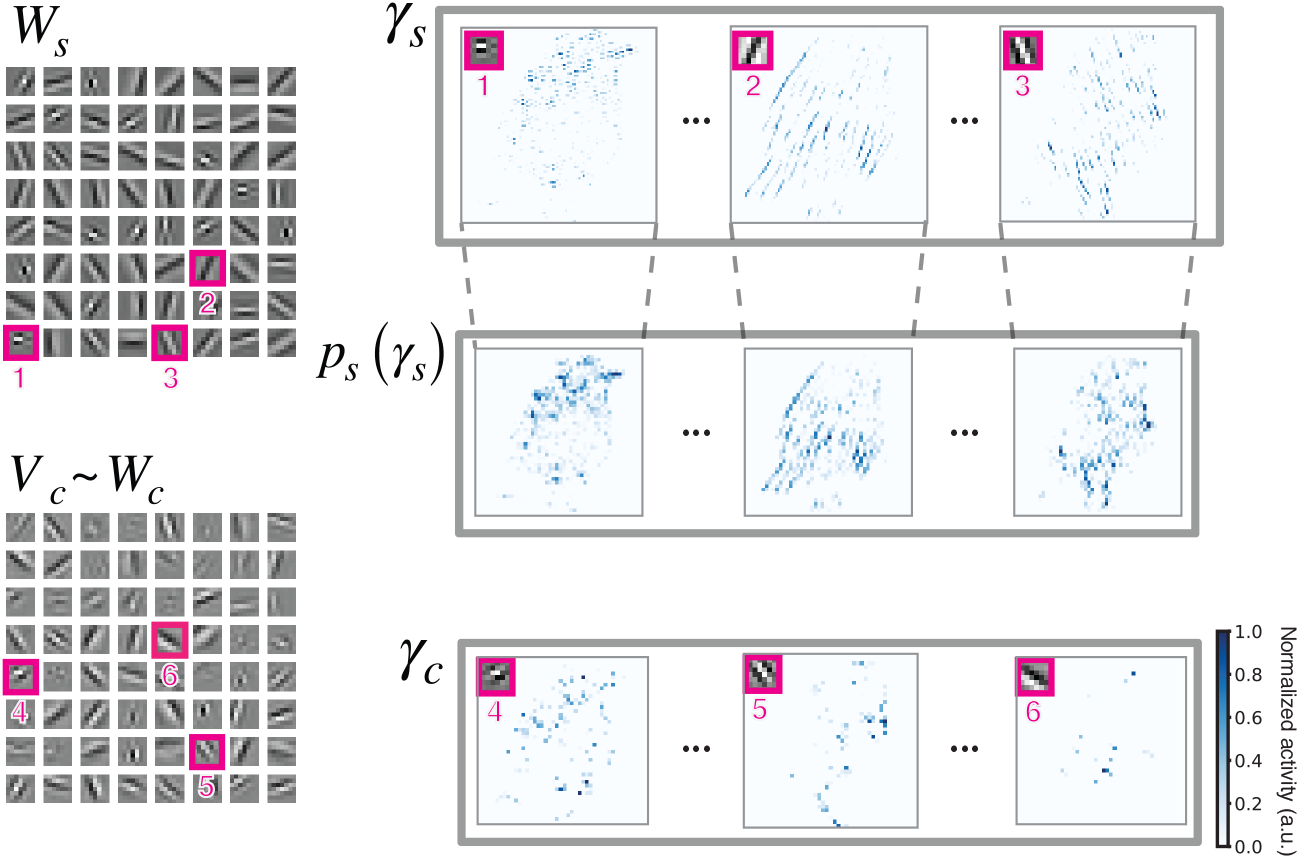
Sparse neural activity maps. **Left.** Example of synaptic weights learned from natural images *W_s_* and *W_c_*, for the first and second layer, respectively. Here we show *V_c_*, a linear approximation of *W_c_* (see Methods 2.8). Each kernel corresponds to a channel in the neural activity maps. **Right.** Representation of 3 channels from the neural activity maps (*γ_s_* and *γ_c_*) elicited by the input *x*. Here, each pixel represents a model neuron and the color code indicates the amplitude of the neural response (lighter for no response and darker blue for the maximal response, here normalized to 1). The kernel in top-left corner indicates the preferred stimulus of each channel.

**Fig S2.**
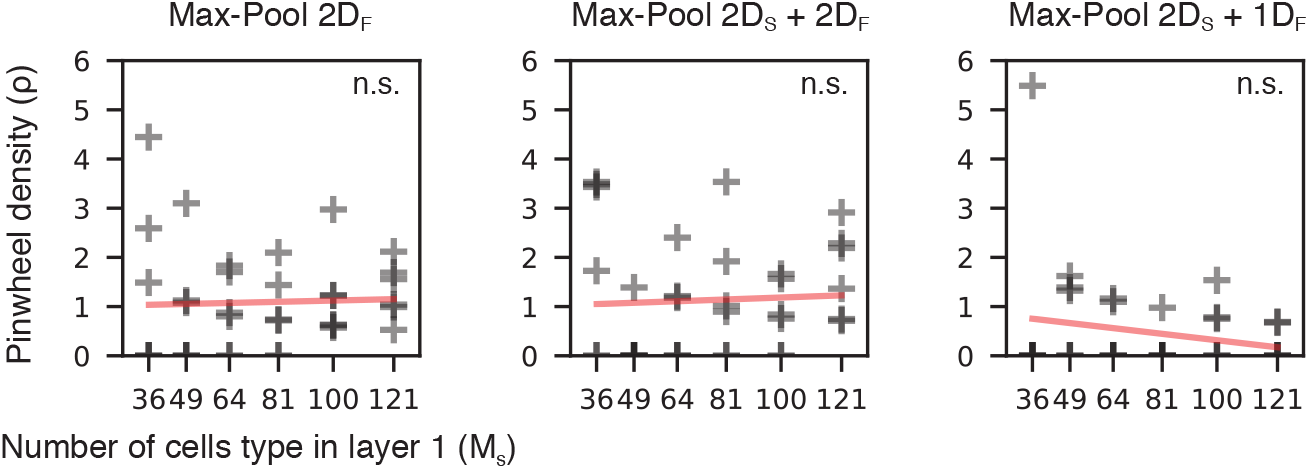
Pinwheels density does not depend on network size. Here we show that there is no relationship between the tested first layer’s network size (*M_s_*) and the density of pinwheels (*ρ*). Pinwheel density is defined as the average number of pinwheel per area unit (see Methods 2.6). For all networks showing orientation maps, we found no significant trend between the size of the map, *M_s_*, and the pinwheel density, in line with neurophysiological observations.

**Fig S3.**
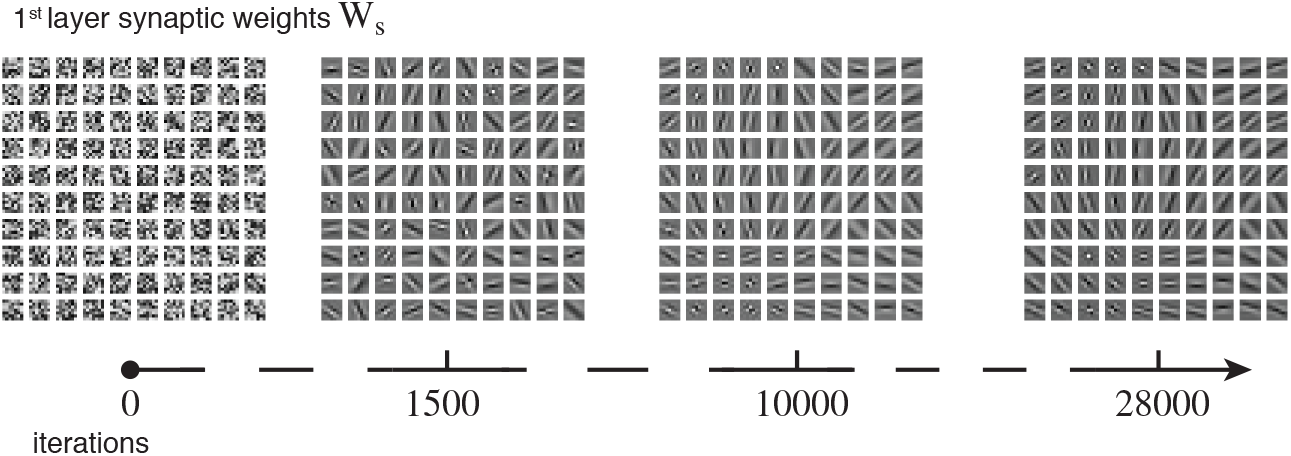
Emergence of topographic maps during learning. We show the evolution of the topographic organization learned by *W_s_* during training when the SDPC network embeds the MaxPool 2*D_S_* + 2*D_F_* function. Here, for *M_s_* = 100. The weights are initialized to random values and the network gradually learns from input data. At first, we observe the emergence of edge detectors similar to the ones observed in the V1 of mammals. Gradually, thanks to the combined action of the feedback coming from the second layer of the network and the pooling function in the forward stream, neighboring cells in the topography become tuned to stimuli of similar orientations but different phases, generating a topographically organized map.

**Fig S4.**
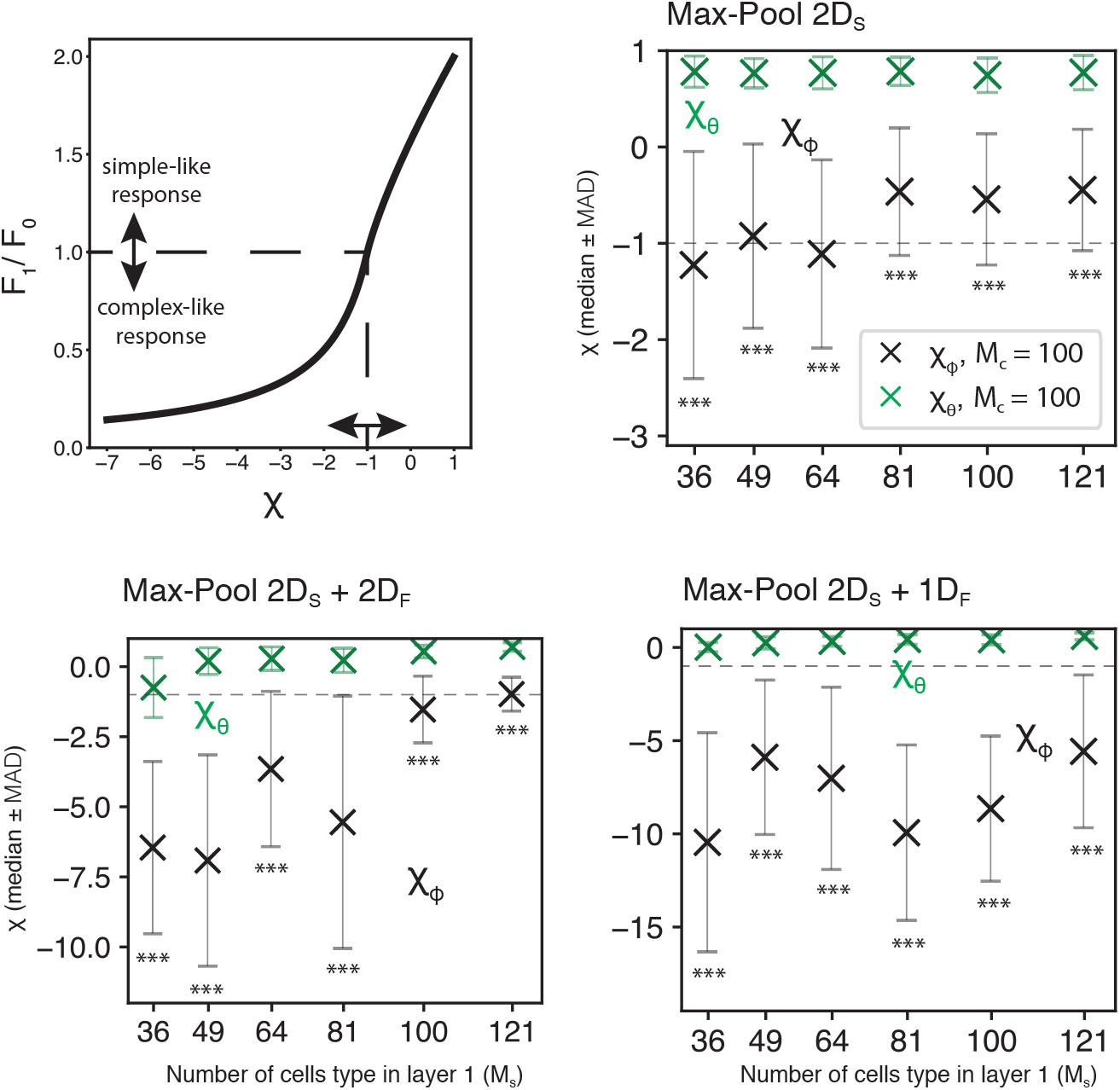
Complex cells population as a function of the network size. We analyze the same results of Fig. 4 in terms of the unimodal variable *χ* as defined in [34] (see, Methods 2.5). In the **top-left** graph we illustrate the nonlinear relationship between *χ* and 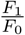. Here the results for *M_c_* = 100 are shown. To avoid the assumption of normally distributed variables, we represent *χ* using the *median ± MAD* (median absolute deviation). Black (*χ_ϕ_*) and green (*χ_θ_*) lines indicate distributions obtained using drifting and rotating grating, respectively. The dashed lines indicate the value *χ* = −1 for which 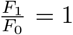, the threshold value for which a V1 cell is typically considered either simple or complex. Using drifting gratings as stimuli significantly generated lower values of *χ_ϕ_* (more complex-like cells) than rotating gratings, in all tested settings (one tailed Wilcoxon signed-rank test). This result confirms that the network’s invariance is linked to the stimulus’ phase and that model cells remain narrowly tuned to orientation. For *p_s_* =MaxPool 2*D_S_* and *p_s_* =MaxPool 2*D_S_* + 1*D_F_* the distribution of *χ_θ_* and *χ_ϕ_* do not vary significantly as a function of the network size. When the network shows a functional topographic map, for *p_s_* =MaxPool 2*D_S_* + 2*D_F_*, we can see a clear dependency between *χ_ϕ_* and the number of features in the first layer of the network *M_s_*.

### S.1.2 Critical points

In this last section, we highlight and discuss some of our model’s critical points and how they could be implemented in a biologically plausible way. First, we wish to state that the main goal of this study is to offer a computational framework able to explain some of the main phenomena observed in mammals’ V1 under a unified computational principle. Here, we are not trying to replicate specific neural circuitry but rather to identify possible mathematical principles that regulate structure and functionalities in biological networks. The SDPC model presented in this study addresses the contribution of three main computational principles: **predictive coding**, **sparse coding** and **pooling**. There is a large set of architectures that could implement such a network in a biological substrate as long as the computation respects simple bio-plausibility rules (e.g., local computation, Hebbian learning, nonnegative firing rates, etc.).

- **Bio-plausibility of max-pooling**: The non-bio-plausible component of max-pooling consists in the fact that the backward network needs to know exactly the index (e.g., the position) of the maximum activity in the previous layer. In other words, the feedback information stream (see Fig. 1) should hold information about which neuron fired with the highest rate for each iteration of the network. This could be easily overcome by approximating the max-pooling with an *ℓ_α_* norm. Indeed, it can be demonstrated that, for any vector *γ*:

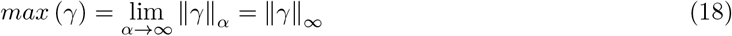 We observed that a norm with a high exponent (*α* = 8) offers qualitatively similar results to the max-pooling. In this case, 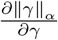 has the advantage of being fully derivable using a local update rule that could be implemented by neural circuitry.
- **Convolution**: In convolution, synaptic weights are repeated across the image to tile the whole visual field, making it impossible to implement a biological system. However, the same retinotopic architecture can be achieved with local untied connections to keep the same structure without replicating the set of weights at each spatial position. [61] showed that such a model converges to a network with the same structure as a convolutional one. In this case, the weights of each plane of a feature map converge to similar configurations, as it would happen in the convolutional case where the weights are forced to be identical. Importantly, the performance is the same as a convolutional network, in terms of classification accuracy, but at the price of a high computational cost. These results suggest that convolutions can be regarded as a good model of retinotopic processing, even if it is not strictly biologically plausible.
- **Symmetry of synaptic weights**: One additional simplification resides in the fact that the SDPC network (and PC networks in general) requires the weights in the recurrent part of the network to be symmetric (see Fig. 1, a). Precisely, the feedback weights in the prediction path correspond to the transpose of those in the feedforward path (respectively 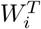 and *W_i_*). The symmetry enforced in the network should not be seen as a significant limitation for the modeling capabilities of the SDPC but, rather, as a simplification needed for computational efficiency. Weight symmetry is a consequence of optimization through gradient descent, but some studies showed that it is unnecessary for the convergence of the network parameters. [62] showed that neural networks can still learn efficient representation even when the feedback weights are fixed to a random distribution. [61] pushed this result even further by showing that not only the two sets of weights can be learned separately by using a Hebbian (local) learning rule but that the weights tend to converge to the transpose of each other (as in the SDPC). This phenomenon is referred to as feedback alignment [63]. Although these last two results were obtained for feedforward neural networks, the derivation of the learning rule for non-symmetric weights is general enough to be easily extended to the SDPC network, as it is exclusively based on partial derivatives and Hebbian learning updates.

## References

1. Hubel DH, Wiesel TN. Receptive Fields, Binocular Interaction and Functional Architecture in the Cat’s Visual Cortex. The Journal of Physiology. 1962;160(1):106–154. doi:10/gdvht8.

2. Hubel DH, Wiesel TN. Receptive Fields and Functional Architecture of Monkey Striate Cortex. The Journal of Physiology. 1968;195(1):215–243. doi:10/gc8z3q.

3. Movshon JA, Thompson ID, Tolhurst DJ. Receptive field organization of complex cells in the cat’s striate cortex. The Journal of physiology. 1978;283(1):79–99.

4. Skottun BC, De Valois RL, Grosof DH, Movshon JA, Albrecht DG, Bonds A. Classifying simple and complex cells on the basis of response modulation. Vision research. 1991;31(7-8):1078–1086.

5. Hubel DH. Brain and visual perception. In: Perception. vol. 34; 2005. p. 3–3.

6. Riesenhuber M, Poggio T. Hierarchical models of object recognition in cortex. Nature neuroscience. 1999;2(11):1019.

7. Mély DA, Serre T. Towards a theory of computation in the visual cortex. In: Computational and cognitive neuroscience of vision. Springer; 2017. p. 59–84.

8. Sakai K, Tanaka S. Spatial pooling in the second-order spatial structure of cortical complex cells. Vision Research. 2000;40(7):855–871. doi:10.1016/S0042-6989(99)00230-8.

9. Antolík J, Bednar JA. Development of Maps of Simple and Complex Cells in the Primary Visual Cortex. Frontiers in Computational Neuroscience. 2011;5(April):1–19. doi:10.3389/fncom.2011.00017.

10. Kim T, Bair W, Pasupathy A. Neural coding for shape and texture in macaque area V4. Journal of Neuroscience. 2019;39(24):4760–4774.

11. Ungerleider LG, Haxby JV. ‘What’and ‘where’in the human brain. Current opinion in neurobiology. 1994;4(2):157–165.

12. Serre T, Oliva A, Poggio T. A feedforward architecture accounts for rapid categorization. Proceedings of the National Academy of Sciences. 2007;104(15):6424–6429. doi:10.1073/pnas.0700622104.

13. Hansen T, Neumann H. A recurrent model of contour integration in primary visual cortex. Journal of Vision. 2008;8(8):8–8.

14. Hoyer PO, Hyvärinen A. A multi-layer sparse coding network learns contour coding from natural images. Vision research. 2002;42(12):1593–1605.

15. Hosoya H, Hyvärinen A. A hierarchical statistical model of natural images explains tuning properties in V2. Journal of Neuroscience. 2015;35(29):10412–10428.

16. Olshausen BA, Field DJ. Sparse coding with an overcomplete basis set: A strategy employed by V1? Vision research. 1997;37(23):3311–3325.

17. Barlow HB, et al. Possible principles underlying the transformation of sensory messages. Sensory communication. 1961;1:217–234.

18. Ringach DL. Spatial structure and symmetry of simple-cell receptive fields in macaque primary visual cortex. Journal of neurophysiology. 2002;88(1):455–463.

19. Rao RP, Ballard DH. Predictive coding in the visual cortex: a functional interpretation of some extra-classical receptive-field effects. Nature neuroscience. 1999;2(1):79.

20. Friston K, Kiebel S. Predictive coding under the free-energy principle. Philosophical Transactions of the Royal Society B: Biological Sciences. 2009;364(1521):1211–1221.

21. Shipp S. Neural elements for predictive coding. Frontiers in psychology. 2016;7:1792.

22. Adesnik H, Naka A. Cracking the Function of Layers in the Sensory Cortex. Neuron. 2018;100(5):1028–1043.

23. Boutin V, Franciosini A, Chavane F, Ruffier F, Perrinet L. Sparse deep predictive coding captures contour integration capabilities of the early visual system. PLoS computational biology. 2021;17(1):e1008629.

24. Boutin V, Franciosini A, Ruffier F, Perrinet L. Effect of top-down connections in Hierarchical Sparse Coding. Neural Computation. 2020;32(11):2279–2309.

25. Carandini M, Heeger DJ. Normalization as a canonical neural computation. Nature Reviews Neuroscience. 2012;13(1):51–62.

26. Tootell RB, Silverman MS, Switkes E, De Valois RL. Deoxyglucose analysis of retinotopic organization in primate striate cortex. Science. 1982;218(4575):902–904.

27. Kaschube M. Neural maps versus salt-and-pepper organization in visual cortex. Current opinion in neurobiology. 2014;24:95–102.

28. Albus K. A quantitative study of the projection area of the central and the paracentral visual field in area 17 of the cat. Experimental Brain Research. 1975;24(2):181–202.

29. Bednar JA, Wilson SP. Cortical maps. The Neuroscientist. 2016;22(6):604–617.

30. Hyvärinen A, Hoyer PO. A two-layer sparse coding model learns simple and complex cell receptive fields and topography from natural images. Vision Research. 2001;41(18):2413–2423. doi:10.1016/S0042-6989(01)00114-6.

31. Hyvärinen A, Hoyer PO, Inki M. Topographic independent component analysis. Neural computation. 2001;13(7):1527–1558.

32. Beck A, Teboulle M. A fast iterative shrinkage-thresholding algorithm for linear inverse problems. SIAM journal on imaging sciences. 2009;2(1):183–202.

33. Sulam J, Papyan V, Romano Y, Elad M. Multi-Layer Convolutional Sparse Modeling: Pursuit and Dictionary Learning. CoRR. 2017;abs/1708.08705.

34. Mechler F, Ringach DL. On the classification of simple and complex cells. Vision research. 2002;42(8):1017–1033.

35. Maass W. On the computational power of winner-take-all. Neural computation. 2000;12(11):2519–2535.

36. LeCun Y, Bengio Y, Hinton G. Deep learning. Nature. 2015;521(7553):436.

37. Coates A, Ng A, Lee H. An analysis of single-layer networks in unsupervised feature learning. In: Proceedings of the fourteenth international conference on artificial intelligence and statistics; 2011. p. 215–12.

38. Wörgötter F, Eysel UT. Quantitative determination of orientational and directional components in the response of visual cortical cells to moving stimuli. Biological cybernetics. 1987;57(6):349–355.

39. Nauhaus I, Benucci A, Carandini M, Ringach DL. Neuronal selectivity and local map structure in visual cortex. Neuron. 2008;57(5):673–679.

40. Fischer S, Šroubek F, Perrinet LU, Redondo R, Cristóbal G. Self-Invertible 2D Log-Gabor Wavelets. International Journal of Computer Vision. 2007;75(2):231–246. doi:10.1007/s11263-006-0026-8.

41. Swindale NV. Orientation tuning curves: empirical description and estimation of parameters. Biological cybernetics. 1998;78(1):45–56.

42. Koch E, Jin J, Alonso JM, Zaidi Q. Functional implications of orientation maps in primary visual cortex. Nature communications. 2016;7(1):1–13.

43. DeAngelis GC, Ghose GM, Ohzawa I, Freeman RD. Functional micro-organization of primary visual cortex: receptive field analysis of nearby neurons. Journal of Neuroscience. 1999;19(10):4046–4064.

44. Kaschube M, Schnabel M, Löwel S, Coppola DM, White LE, Wolf F. Universality in the evolution of orientation columns in the visual cortex. science. 2010;330(6007):1113–1116.

45. Karklin Y, Lewicki MS. Emergence of complex cell properties by learning to generalize in natural scenes. Nature. 2009;457(7225):83–86. doi:10.1038/nature07481.

46. Stevens JLR, Law JS, Antolík J, Bednar JA. Mechanisms for stable, robust, and adaptive development of orientation maps in the primary visual cortex. Journal of Neuroscience. 2013;33(40):15747–15766.

47. Swindale NV, Shoham D, Grinvald A, Bonhoeffer T, Hübener M. Visual cortex maps are optimized for uniform coverage. Nature neuroscience. 2000;3(8):822–826.

48. Heimel JA, Van Hooser SD, Nelson SB. Laminar organization of response properties in primary visual cortex of the gray squirrel (Sciurus carolinensis). Journal of neurophysiology. 2005;94(5):3538–3554.

49. Murphy EH, Berman N. The rabbit and the cat: a comparison of some features of response properties of single cells in the primary visual cortex. Journal of Comparative Neurology. 1979;188(3):401–427.

50. Gilbert CD. Laminar differences in receptive field properties of cells in cat primary visual cortex. The Journal of physiology. 1977;268(2):391–421.

51. Schiller PH, Finlay BL, Volman SF. Quantitative studies of single-cell properties in monkey striate cortex. I. Spatiotemporal organization of receptive fields. Journal of neurophysiology. 1976;39(6):1288–1319.

52. Martinez LM, Alonso JM. Complex receptive fields in primary visual cortex. The neuroscientist. 2003;9(5):317–331.

53. Bereshpolova Y, Hei X, Alonso JM, Swadlow HA. Three rules govern thalamocortical connectivity of fast-spike inhibitory interneurons in the visual cortex. Elife. 2020;9:e60102.

54. Ohzawa I, DeANGELIS GC, Freeman RD. Encoding of binocular disparity by complex cells in the cat’s visual cortex. Journal of neurophysiology. 1997;77(6):2879–2909.

55. Fleet DJ, Wagner H, Heeger DJ. Neural encoding of binocular disparity: energy models, position shifts and phase shifts. Vision research. 1996;36(12):1839–1857.

56. Van den Bergh G, Zhang B, Arckens L, Chino YM. Receptive-field properties of V1 and V2 neurons in mice and macaque monkeys. Journal of Comparative Neurology. 2010;518(11):2051–2070.

57. Hansel D, van Vreeswijk C. The mechanism of orientation selectivity in primary visual cortex without a functional map. Journal of Neuroscience. 2012;32(12):4049–4064.

58. Van Hooser SD. Similarity and diversity in visual cortex: is there a unifying theory of cortical computation? The Neuroscientist. 2007;13(6):639–656.

59. Niell CM, Stryker MP. Highly selective receptive fields in mouse visual cortex. Journal of Neuroscience. 2008;28(30):7520–7536.

60. Jang J, Song M, Paik SB. Retino-Cortical Mapping Ratio Predicts Columnar and Salt-and-Pepper Organization in Mammalian Visual Cortex. Cell Reports. 2020;30(10):3270–3279.

61. Amit Y. Deep learning with asymmetric connections and Hebbian updates. Frontiers in computational neuroscience. 2019;13:18.

62. Lillicrap TP, Cownden D, Tweed DB, Akerman CJ. Random synaptic feedback weights support error backpropagation for deep learning. Nature communications. 2016;7(1):1–10.

63. Lillicrap TP, Santoro A, Marris L, Akerman CJ, Hinton G. Backpropagation and the brain. Nature Reviews Neuroscience. 2020; p. 1–12.

